# Scalable cell-free production of active T7 RNA polymerase

**DOI:** 10.1101/2024.11.19.623967

**Authors:** Ryan N. Rezvani, Rochelle Aw, Wei Chan, Krishnathreya Satish, Han Chen, Adi Lavy, Swechha Rimal, Divyesh A. Patel, Erik Kvam, Ashty S. Karim, Antje Krüger, Weston K. Kightlinger, Michael C. Jewett

**Affiliations:** Cell-free Protein Synthesis and Microbial Process Development, National Resilience, Inc, 3115 Merryfield Row, San Diego, California 92121, United States; Department of Bioengineering, Stanford University, Stanford, California 94305, United States; GE HealthCare Technology & Innovation Center, Niskayuna, New York 12309, United States; Department of Chemical and Biological Engineering, Northwestern University, Evanston, Illinois 60208, United States

**Keywords:** Cell-free gene expression, in-vitro transcription and translation, T7 RNA polymerase, scale-up, biomanufacturing

## Abstract

The SARS-CoV-2 pandemic highlighted the urgent need for biomanufacturing paradigms that are robust and fast. Here, we demonstrate the rapid process development and scalable cell-free production of T7 RNA polymerase, a critical component in mRNA vaccine synthesis. We carry out a one-liter cell-free gene expression (CFE) reaction that achieves over 80% purity, low endotoxin levels, and enhanced activity relative to commercial T7 RNA polymerase. To achieve this demonstration, we implement rolling circle amplification to circumvent difficulties in DNA template generation, and tune cell-free reaction conditions, such as temperature, additives, purification tags and agitation to boost yields. We achieve production of a similar quality and titer of T7 RNA polymerase over more than 4 orders of magnitude reaction volume. This proof of principle positions CFE as a viable solution for decentralized biotherapeutic manufacturing, enhancing preparedness for future public health crises or emergent threats.

## Introduction

Cell-based biomanufacturing practices for therapeutics have several limitations. For example, the timeline for developing protein therapeutics is long (typically >18 months). In addition, cell lines must be developed, selected, maintained, and expanded anew for each protein biologic (i.e., one bug, one drug paradigm). Moreover, current manufacturing approaches require significant infrastructure to meet demand (Mao & Chao, 2020). These challenges were highlighted during the SARS-CoV-2 pandemic where speed and consistency were fundamental to the production of high-quality vaccines (Rosa et al., 2021). Suppliers were unable to meet demands of some raw material components like T7 RNA polymerase (T7 RNAP), a critical enzyme for synthesizing mRNA vaccines which is produced by centralized, cellular manufacturing methods and accounts for ∼ 34% of the overall raw material cost (Guan et al., 2020; Raghuwanshi et al., 2024). While mRNA vaccines are an attractive and rapid alternative manufacturing paradigm to conventional vaccines (Jackson et al., 2020), this shift to “cell-free” manufacturing principles does not yet go beyond vaccines. Developing rapid manufacturing approaches for enzymes and therapeutic proteins will be essential to effectively address future public health emergencies and growing demands in personalized healthcare.

Cell-free gene expression (CFE) offers a promising alternative to cellular manufacturing, creating an opportunity for decentralized production (Borhani et al., 2023; Bundy et al., 2018; Hunt, 2024; Lee & Kim, 2024; Pardee et al., 2016; Silverman, Karim, et al., 2020; Warfel et al., 2023). CFE has offered speed and control for a number of different applications including enzyme prototyping (Ekas, 2024a; Karim et al., 2020; Karim & Jewett, 2016; Kightlinger et al., 2019; Kightlinger et al., 2018; Liew et al., 2022; Vögeli et al., 2022), biosensor development (Ekas, 2024b; Nguyen et al., 2021; Sadat Mousavi et al., 2020; Silverman, Akova, et al., 2020; Thavarajah et al., 2020), studying protein-protein interactions (Hunt et al., 2023), building artificial cells (Peruzzi et al., 2023), and education (Collins et al., 2024; Huang et al., 2018; Stark et al., 2019; Stark et al., 2018), among others. However, CFE can go beyond prototyping to produce therapeutically relevant products. Work at laboratory scale using CFE to produce effective therapeutics have included producing virus-like particles (Chan et al., 2015; Spice et al., 2020), antiviral proteins (Borhani et al., 2023), and vaccines (Stark et al., 2021; Williams et al., 2023). Despite these advancements, the challenge of scaling CFE to meet urgent manufacturing needs remains. While CFE has demonstrated versatility and efficiency in laboratory settings, limited examples of scale beyond milliliter volumes have been demonstrated. While notable exceptions, such as the production of cytokines and antibodies illustrate the potential of CFE for industrial production (Yin et al., 2012; Zawada et al., 2011), the use of the technology ‘at scale’ is not widely accessible, and CFE is yet to be implemented to produce mRNA manufacturing enzymes or in decentralized settings. Overall, the need for rapid production methods for therapeutics, vaccines, and mRNA manufacturing enzymes remains.

Here, we set out to transform protein manufacturing by developing a process to produce a medically-relevant enzyme within one month at scale using CFE. As a model system, we selected the production of T7 RNAP, a protein projected to increase in demand due to therapeutic advances in vaccines (Qin et al., 2022). Our design criteria included that the CFE-produced T7 RNAP had to (i) have purity of >80%, (ii) be produced at yields of >0.1 g/L, (iii) have low endotoxin levels, below 12 EU per 10 µg of protein, and (iv) show activity similar to a commercially available T7 RNAP. Our findings support the emergence of distributed manufacturing approaches for medical products, one that leverages the unique advantages of CFE to meet urgent healthcare needs.

## Materials and Methods

### DNA preparation

Plasmid DNA was prepared using the QIAGEN Plasmid Midi Kit (QIAGEN, Germantown, Maryland, USA) according to the manufacturer’s protocols. Gibson assembly reactions were performed to generate linear expression templates and rolling circle amplification (RCA) product for CFE. For this, pJL1 backbone was ordered as gBlock from Integrated DNA Technologies, Inc (IDT; Coralville, Iowa, USA). sfGFP and T7 RNAP to be produced by CFE were codon-optimized using the IDT codon optimization tool and ordered from IDT as gBlocks containing pJL1 Gibson assembly overhangs. Gibson assembly was used to assemble sfGFP or T7 RNAP open reading frame DNA with the pJL1 backbone following published protocols (Gibson et al., 2009). Linear expression templates (LETs) were prepared from Gibson assembly of T7 RNAP with pJL1-plasmid backbone, diluting the completed reaction 20-fold, and subsequent PCR using Q5 Hot Start DNA polymerase (New England Biolabs, Ipswich, Massachusetts, USA) and pJL1 LET primers (ctgagatacctacagcgtgagc, cgtcactcatggtgatttctcacttg (Hunt et al., 2023; Krüger et al., 2020)) following manufacturer’s instructions. RCA reactions were prepared using 20-fold diluted completed Gibson assembly reaction, EquiPhi29™ DNA Polymerase (Thermo Fisher Scientific, Waltham, Massachusetts), and exo-resistant primers binding to pJL1-plasmid backbone according to the manufacturer’s protocol if not stated otherwise. Optimized RCA conditions refer to following the manufacturer’s protocol but using 1.5-fold dNTPs and 0.25-fold EquiPhi29™ DNA Polymerase.

### Cell extract preparation

Cell extract was prepared from BL21 star (DE3). The strain choice of BL21 star (DE3) was chosen due to observed cleavage of RNA polymerase, particularly those that lack an OmpT deletion (Grodberg & Dunn, 1988; Muller et al., 1988). BL21 star (DE3) was grown in rich media supplemented with additional carbon source according to Resilience’s proprietary protocol. Cultures were grown to an OD600 of 5 before induction with 1 mM of isopropyl β-d-1-thiogalactopyranoside and grown to a final OD600 of 37. Cells were pelleted at 7,100 *x g* for 10 minutes at 4ºC and washed twice with cold S30 buffer (14 mM magnesium acetate, 60 mM potassium acetate, 10 mM Tris base pH 8.2), resuspended with S30 buffer, and lysed using an Avestin EmulsiFlex B15 at 20,000 psi using a single pass. The resulting lysate was clarified twice at 18,000 *x*g for 20 minutes at 4ºC, flash-frozen, and stored at -80ºC until further use.

### CFE reactions

CFE reactions were run using BL21 star (DE3) extracts and a proprietary reagent system. In small-scale reactions (15 µL), CFE optimization experiments were performed. FkpA was tested in concentrations ranging from 0-75 µM, DTT was tested from 0-16 mM, and RCA product was added in dilutions ranging from 250-20,000-fold. Small scale reactions were performed at 15 µL scale in 2 mL Eppendorf tubes and left to incubate at 25 or 30ºC for 22 hours. To visualize T7 RNAP expression via SDS-PAGE, proteins were labeled during CFE with FluoroTect™ (Promega, Madison, Wisconsin, USA). FluoroTect™ was added to CFE reactions at 3.33% v/v. For 30 mL CFE reactions, square bioassay dishes (Corning) were used and placed in an incubator at 25ºC with 250 rpm shaking. The reactions were incubated for 16-22 hours. Used CFE conditions at 30 ml scale were as described above but without FkpA, 8 mM DTT, and 15 µl of 1200-fold diluted RCA product. The 1-L CFPS were performed in a 1 L bioreactor using 500 µL of RCA product. The reactions were performed at 25ºC with DO controlled by agitation and airflow.

### Protein purification

CFE samples were centrifuged at 16,000 *xg* for 15 minutes at 4ºC and the supernatant transferred into a fresh 15 mL conical tube. His-tagged proteins were purified using a 1 ml HisTrap Excel resin column (Cytiva, Wilmington Delaware, USA) with an ÄKTA Avant (Cytiva) according to manufacturer’s protocols. Briefly, the column was equilibrated with 10 CV of Buffer A (20 mM sodium phosphate pH 7.4, 500 mM sodium chloride) containing 10 mM imidazole. The clarified CFE reaction was diluted 2-fold with Buffer A containing 10 mM imidazole and loaded on to the column. The column then was washed with 10 CV of Buffer A plus 10% Buffer B (20 mM sodium phosphate pH 7.4, 500 mM sodium chloride, 500 mM imidazole). His-tagged T7RNAP was eluted with 4-100% Buffer B over 10 CV and 10 CV Buffer B strip. Elution fractions were collected at 4°C in 96-deep well plates.

Strep-tagged proteins were purified using a 1 mL Strep-tactin XT Sepharose column (Cytiva) with an ÄKTA Avant (Cytiva) according to manufacturer’s protocols. Briefly, the column was equilibrated with 10 CV on Buffer A (PBS, Corning, Glendale, Arizona, USA) before 30 mL of the clarified CFE reaction was loaded on to the column followed by 10 mL of Buffer A. To wash the column 10 CV of Buffer A was added, and elution was performed using a gradient from 0 to 100% using Buffer B (PBS, 50 mM biotin; IBA Life sciences, Göttingen, Germany). Elution fractions were collected at 4°C in 96-deep well plates (Cytiva).

### Endotoxin removal

Endotoxins were removed using Pierce™ High Capacity Endotoxin Removal Resin (Thermo Fisher Scientific). All buffers were prepared using endotoxin-free water (Sigma Aldrich, St Louis, Missouri, USA). Prior to use 2 mL of resin was packed into a 15 mL conical tube and washed with 10 mL of 0.2 N sodium hydroxide and incubated for 10 minutes before being removed using a pipette preventing disruption of the resin bed. A further 10 mL of 10 N sodium hydroxide was added and mixed before incubation at room temperature for 14-16 hours. The liquid was then removed using a pipette, and 10 mL of 2M sodium chloride was added, incubated for 10 minutes and then removed. This step was repeated, before 10 mL of endotoxin free water was added, incubated for 10 minutes and repeated for a final time. Finally, 1 mL of endotoxin free water was added to the resin to create the resin slurry prior to being added to the column.

The regenerated resin was added to a 2 mL Poly-Prep® Chromatography column (Bio-Rad Laboratories, Hercules, California, USA). The resin was equilibrated with 1 mL of PBS and repeated a further three times for a total of 4 mL. The pooled purified elution fractions were diluted two-fold in PBS and was added to the resin in 1 mL aliquots.

### Diafiltration

Post-endotoxin removal samples were loaded on to an Amicon Ultra-15 centrifuge filter MWCO 10 KDa (Sigmal Aldrich). The samples were concentrated at 5000 *x*g for 15 minutes at 4°C. The retentate should concentrate to 1 mL, if necessary, centrifuge at 5000 *x*g for 10 minutes at 4°C until the appropriate volume is reached. The retentate is diluted 10-fold using the formulation buffer (20 mM Tris, 150 mM sodium chloride, 0.5 mM DTT, pH 8, made with endotoxin free water). Samples were concentrated to ∼1.25 mL and passed through a 0.22 µm PES syringe filter using a 5 mL syringe.

### Endotoxin assay

Endotoxin concentrations were measured either using the Endosafe® nexgen-PTS™ system (Charles River Laboratories, Wilmington, Massachusetts. USA) or the Pierce™ Chromogenic Endotoxin Quant Kit (Thermo Fisher Scientific) according to the manufacturers’ protocols.

### Protein analysis

Samples were prepared by heating to 70ºC for 3 minutes in the Invitrogen™ NuPAGE™ LDS Sample Buffer (Thermo Fisher Scientific) and loaded on a 4-12% Bis-Tris gel and run with MOPS SDS running buffer at 160V until the dye reached the end of the gel. The gels were then stained with InstantBlue (AbCam, Waltham, Massachusetts, USA) for 1 hour shaking at room temperature, and then destained with milliQ water for 1 hour shaking at room temperature. Gels were either imaged using an Amersham Typhoon (Cytiva) and channel setting [IRShort] or an Amersham Image Quant 800 and channel setting OD600 (Cytiva).

### T7 RNA polymerase assay

To determine the relative activity of CFE-produced T7 RNAP to a commercially available T7 RNAP standard (GenScript Biotech, Piscataway, New Jersey, USA), the T7 RNAP assay was performed using the T7 RNAP Polymerase Assay Kit according to manufacturer’s protocols (Profoldin, Hudson, Massachusetts, USA; Catalog number T7RPA100K).

## Results

### Overcoming self-activation of the T7 RNA promoter using rolling circle amplification

We first set out to optimize a CFE DNA template for high expression of T7 RNAP. Standard DNA templates used in CFE are T7 promoter driven plasmids that lack repressor regulatory elements (i.e., devoid of lac operator sequence) to maximize protein expression (Jew et al., 2022). When we attempted to clone and propagate the T7 RNAP plasmid by bacterial fermentation, we were not successful. We found that the plasmid encoding T7 RNAP became mutated. This is likely because leaky expression of T7 RNAP triggered a positive feedback loop with the T7 promoter that created excessive burden on the system (Chen et al., 1994). Therefore, obtaining plasmid-based DNA for large volume CFE reactions was not feasible.

Synthetic linear expression templates (LETs) that do not require propagation in *Escherichia coli* work well for CFE expression (McSweeney & Styczynski, 2021; Sun et al., 2014). We constructed LETs with a T7 promoter and terminator by Gibson assembly of T7 RNAP open reading frame into pJL1-plasmid backbone and subsequent PCR. We then supplemented these LETs at varied concentrations in CFE reactions to produce T7 RNAP, which was measured by incorporation of a fluorescent amino acid (**Figure 1A**). Crude extracts were derived from BL21 star (DE3) cells. These cells lack OmpT, which is known to cleave T7 RNAP (Davanloo et al., 1984; Tabor & Richardson, 1985). LETs resulted in poor expression of the T7 RNAP, which may be because linear DNA is known to be unstable in extracts due to nuclease sensitivity (Chen & Lu, 2021).

**Figure 1.**
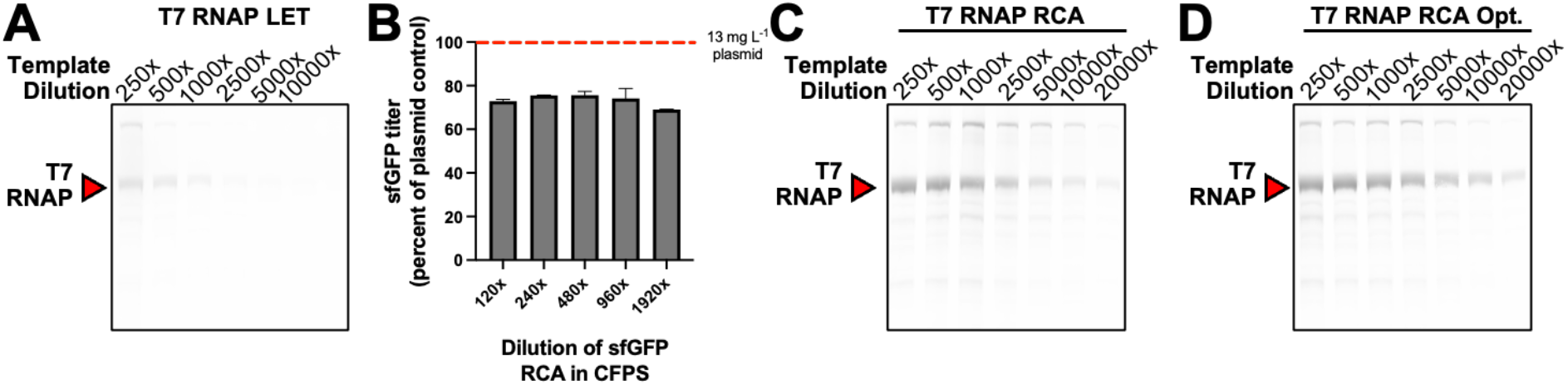
Overcoming self-activation of the T7 RNA promoter using rolling circle amplification. A) SDS-PAGE showing FluoroTect™ selective detection of soluble, CFE-synthesized protein produced either from a linear expression template (LET). B) Durability of sfGFP CFE titers with decreasing amounts of RCA template. sfGFP productivity is written as a percentage of the standard pDNA control (in gray). C) SDS-PAGE showing FluoroTect™ selective detection of soluble, CFE-synthesized protein produced either from rolling circle amplification (RCA). D) SDS-PAGE showing FluoroTect™ selective detection of soluble, CFE-synthesized protein produced either from optimized rolling circle amplification recipe (RCA Opt.) Gels are representative of n=3 independent experiments.

To scale DNA template preparation and improve expression yields, we next tried to prepare CFE templates by using rolling circle amplification (RCA), an isothermal enzymatic synthesis reaction involving ø29 DNA polymerase (Lizardi et al., 1998). RCA has previously been used for CFE applications (Dopp et al., 2019; Hadi et al., 2020; Kumar & Chernaya, 2009). First, we tested the suitability of RCA product in our CFE system for producing sfGFP by setting up RCA reactions using Gibson assembly reactions of pJL1 backbone with sfGFP gBlocks as template, exo-resistant primers, and EquiPhi29™ DNA Polymerase. We supplemented RCA product at varied concentrations in CFE reactions and compared sfGFP titers to that produced from a standard plasmid-based CFE reaction at 13 mg L^-1^ of plasmid DNA (**Figure 1B**). The RCA template resulted in ∼25% lower sfGFP yield of protein compared to plasmid DNA. Of note, in contrast to T7 RNAP plasmid, sfGFP plasmid did not have issues being prepared from cells. As plasmid DNA components account for one of the highest costs of a CFE system (Carlson et al., 2012; Silverman, Karim, et al., 2020; Stark et al., 2023), using RCA products lowers the cost of CFE reactions (Hadi et al., 2020).

Next, we created T7 RNAP expression templates using RCA and supplemented them at varied concentrations in CFE reactions (diluting the template up to 20,000 times). We measured T7 RNAP production from the RCA products by incorporating FluoroTect™ lysine amino acid into the CFE reaction, determined by an SDS PAGE gel (**Figure 1C**). The diluted RCA template was able to support robust protein expression, maintaining ∼50% expression titers with up to a 2,500-fold dilution. To achieve higher yields, while reducing costs per RCA reaction, we tested T7 RNAP production using an optimized RCA recipe that contained 1.5x NTPs and 0.25x of EquiPhi29™ DNA Polymerase. Given equivalent starting RCA templates, we found that higher yields were achieved with this optimized RCA recipe in comparison to the non-optimized version (**Figure 1D**). The ability to use RCA products reduces the protein production timeline by removing the necessity to propagate DNA in *E. coli*, further improving the ability to respond to a novel threat at speed.

### Optimizing expression of T7 RNA polymerase

We next sought to increase the production of T7 RNAP by altering the CFE reaction conditions. Lowering the expression temperature can improve yield of hard to express proteins in CFE (Chou, 2007; Jin et al., 2019). We compared expression of T7 RNAP at 30 ºC and 25 ºC (**Figure 2A**). From a visual inspection of the expression levels of T7 RNAP on gels between the two temperatures, we observed a modest improvement when expression occurs at a lower temperature. As a result, this condition was set for further experiments.

**Figure 2.**
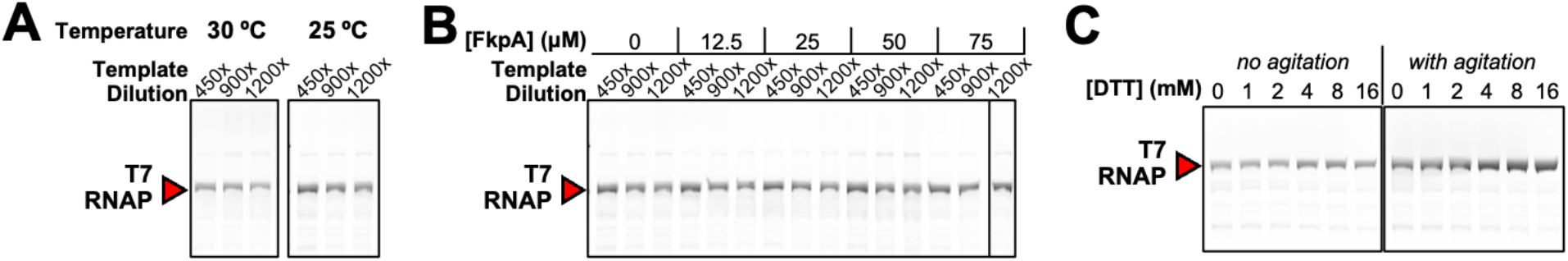
Optimizing expression of T7 RNA polymerase. A) SDS-PAGE showing FluoroTect™ selective detection of soluble, CFE-synthesized protein produced at either 30°C or 25°C. B) CFE reaction optimizations for T7 RNAP at 25°C, ran on SDS-PAGE and selectively imaged with FluoroTect™. [FkpA] refers to the concentration of supplemented *E. coli* prolyl isomerase. C) CFE reaction optimizations for T7 RNAP, ran on SDS-PAGE and selectively imaged with FluoroTect™ at 25°C. [DTT] refers to the concentration of the reducing agent dithiothreitol with and without agitation. Gels are representative of n=3 independent experiments.

While T7 RNAP does not contain any disulfide bonds, the addition of peptidyl-prolyl cis-trans isomerases may have benefits for improving folding kinetics and stabilizing the protein (Ramm & Plückthun, 2000). We thus added the chaperone FKBP-type peptidyl-prolyl cis-trans isomerase (FkpA) to the CFE reaction to determine its effect on the production of T7 RNAP (**Figure 2B**). At 25 ºC, we did not see a difference in yield with FkpA. However, we do see slight increase when expressed at 30 ºC (**Supplementary Figure 1**), which may be expected as lower temperatures often result in improved folding of complex proteins (Chou, 2007). Therefore, higher temperatures have more propensity of misfolding, which the FkpA additive may help to prevent. As FkpA showed an improvement at the higher temperatures, and no detrimental impact at the lower temperatures, FkpA was included in all future experiments.

Oxygen availability has been shown to be a limiting factor in CFE reactions (Jewett et al., 2008) and in fed batch conditions could be exacerbated. Therefore, we next evaluated the impact of using agitation to increase oxygen transfer (**Figure 2C**). Shaking at 250 rpm improves the production of T7 RNAP compared to a non-shaking condition. Simultaneously, we determined whether the addition of dithiothreitol (DTT) to create a reduced environment was beneficial to the production of T7 RNAP (**Figure 2C**). Both with and without agitation, DTT boosted protein yield by 33%, and this effect was even more pronounced with agitation (70% protein yield increase compared to the no DTT condition).

### Optimizing purification of T7 RNA polymerase

To utilize the T7 RNAP downstream purification was optimized to achieve protein samples that were over 80% pure with endotoxin levels within the regulated level of <12 EU per 10 µg of protein. CFE reactions were carried out in large petri dishes at the 30mL scale. A three-step purification process was employed, the first being affinity chromatography followed by a column-based endotoxin removal step, and finally the samples were buffer exchanged into an appropriate storage buffer (**Figure 3A**). Two separate purification tags were evaluated, a polyhistidine (His)-tag (HHHHHH) (**Supplementary Figure 2A**) and a strep-tag (WSHPQFEK) (**Supplementary Figure 2B**). We measured the relative yields from a Ni-Sepharose or Strep-Tactin purification that had undergone endotoxin removal using the Pierce™ High Capacity Endotoxin Removal Resin (**Figure 3B**). We observed 0.13 g/L in yields. Purity was calculated by densitometry, with both purification tags exceeding our initial 80% benchmark for purity. The yields obtained using the two different tags were similar, but the strep-tactin resin has a higher binding capacity, which is advantageous for scaling up the process.

**Figure 3.**
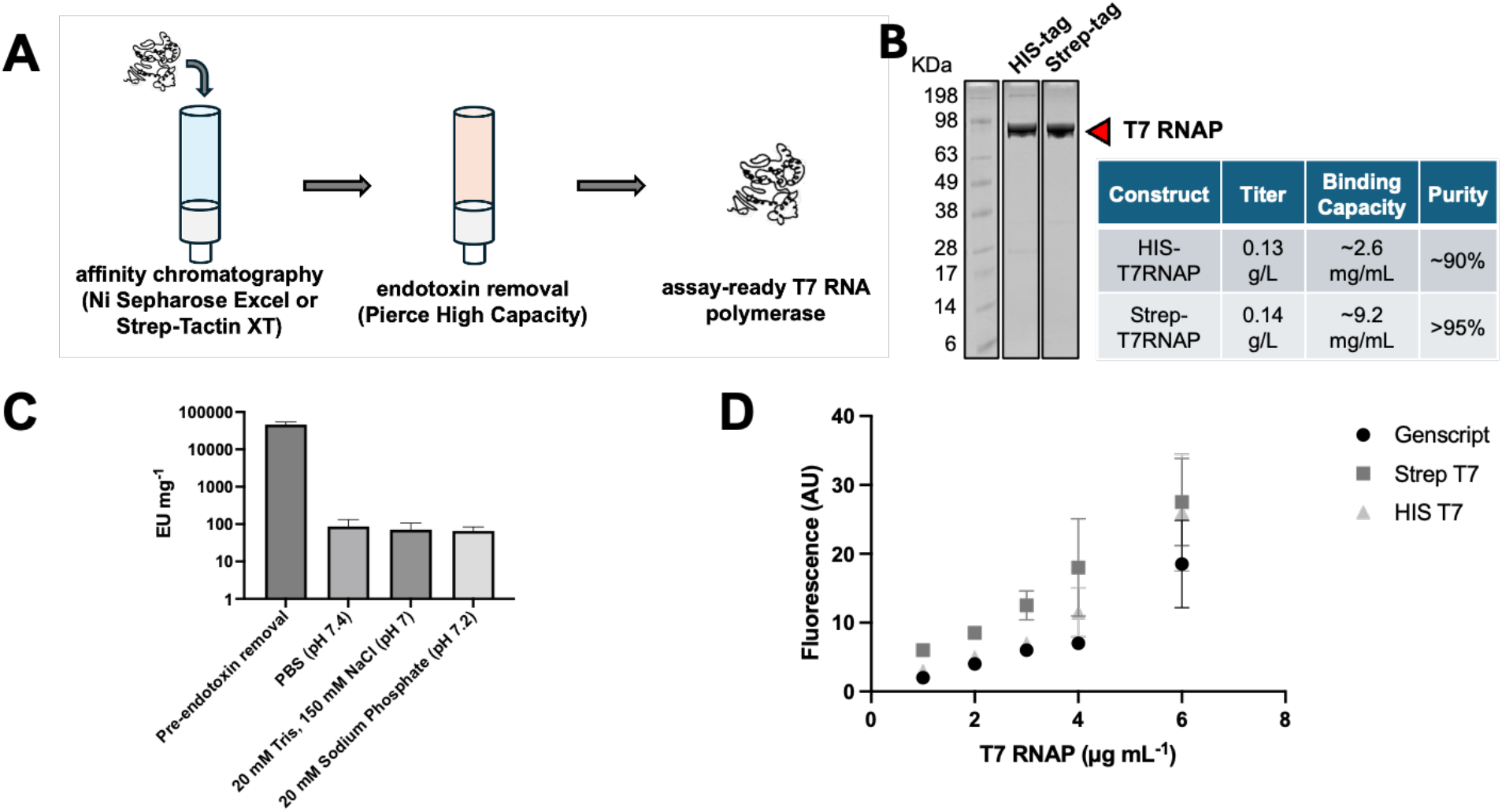
Optimizing purification of T7 RNA polymerase. A) Schematic showing 2-step chromatography workflow to produce low-endotoxin, assay-ready enzyme. B) Coomassie-stained SDS-PAGE depicting the purified, endotoxin-removed, and buffer-exchanged T7 RNAP from either the Ni Sepharose (His-tag) or Strep-Tactin (Strep-tag) processes. Table on the right details the titer of each construct (per liter of CFE reaction) and the binding capacity (per mL of affinity resin). C) Endotoxin removal of Strep-Tactin XT-purified T7 RNAP with various equilibration buffers. D) Fluorescence-based activity of the HIS-tagged and Strep-tagged T7 RNAP produced in 30 mL petri-dish CFE reactions, as compared to the commercially available T7 RNAP from Genscript. Data are representative of n=3 independent experiments, with standard deviation shown.

To determine the efficiency of the endotoxin removal, samples were quantified using the Endosafe® nexgen-PTS™ system (**Figure 3C**). Endotoxin levels decreased over 1000-fold pre-endotoxin removal and were tested in various formulation buffers. All three formulations buffers showed similar endotoxin levels at 0.869 EU per 10 µg for PBS, 0.709 EU per 10 µg for 20 mM Tris, 150 mM NaCl (pH 7), and 0.662 EU per 10 µg for 20 mM sodium phosphate (pH 7.2). All samples showed endotoxin levels below the positive control from Genscript and the threshold we set of 12 EU per 10 µg, which is a standard in therapeutic conjugate vaccines (Bolgiano et al., 2007; Brito & Singh, 2011). The final formulation for purification was 20 mM Tris, 150 mM NaCl, 0.5 mM DTT, pH 8, and was chosen for improved stability of the T7 RNAP.

We then tested the activity of the T7 RNAP was tested in comparison to a commercial T7 RNAP (**Figure 3D**). T7 RNAP with a strep tag or with a his-tag resulted in improved activity compared to the commercial version. At the highest concentration of 4 µg mL^-1^ the activity difference was 148% for the strep-tag purified T7 RNAP, and 140% or the his-tag purified T7 RNAP.

### Tech transfer shows reproducibility

Interlaboratory variability is a known limitation for wider adoption of CFE systems (Cole et al., 2019). This variability goes beyond the production of difficult to produce components such as the extract and reagents, to also include operator, equipment and day-to-day variability. To determine if an alternative site could produce similar quantities and qualities of T7 RNAP, extract and reagents were sent to a second site and reactions, purifications, and further downstream processing were set up by a different operator.

CFE reactions were set up at a 30 mL scale in petri dishes, using the strep-tag construct. Samples were purified and endotoxin was removed using the optimized expression and purification strategy. Yields of protein were lower than had previously been achieved at 0.034 mg mL^-1^ compared to 0.12 mg mL^-1^ during process development. These differences may be due to differences in equipment used as excessive evaporation occurred during the CFE reaction that was not previously reported.

The diafiltrated samples were run on an SDS-PAGE gel and showed 84.9% purity (**Figure 4A**), which is lower than what was observed at the original test, but above the metric of 80% purity. Endotoxin levels were measured using the Pierce™ Chromogenic Endotoxin Quant Kit and resulted in final endotoxin levels of 1.12 EU per 10 µg of protein, which is under the 12 EU per 10 µg target (**Figure 4B**). Finally, activity tests were performed and had been observed previously, the purified sample outperformed the state of the art purchased control showing 125% of activity (**Figure 4C**). While the quality of the product did not match identically to those produced at the primary site, the ability to transfer the technology to another lab with only a written protocol for guidance shows the adoptability of CFE and the potential to use this technology for decentralized manufacturing and multiple sites worldwide.

**Figure 4.**
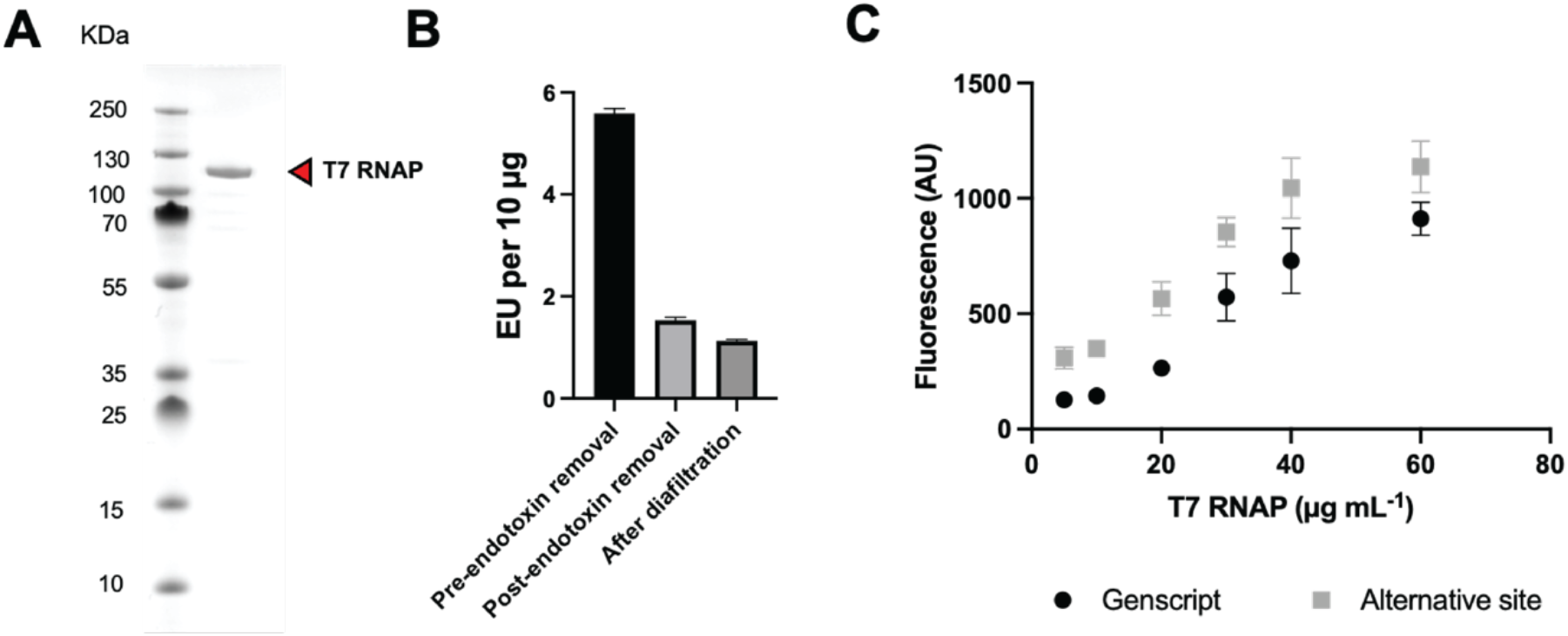
Reproducible expression, purification, and activity of T7 RNA polymerase across laboratories. A) Coomassie-stained SDS-PAGE depicting the purified, endotoxin-removed, and buffer-exchanged T7 RNAP. B) Endotoxin removal of Strep-Tactin XT-purified T7 RNAP pre-removal, post-removal and post-diafiltration. C) Fluorescence-based activity of the HIS-tagged and Strep-tagged T7 RNAP produced in 30 mL petri-dish CFE reactions, as compared to the commercially available T7 RNAP from Genscript. Data are representative of n=3 independent experiments, with standard deviation shown.

### Scale-up at the 1-L scale

To show the potential of large-scale production in CFE, our process was scaled to a 1-L bioreactor. To validate the bioreactor run, a side-by-side comparison in 30 mL reactions in petri dishes was run concurrently. An SDS-PAGE gel shows equivalent accumulation of the T7 RNAP over the 22-hour incubation (**Figure 5A**). Downstream processing was performed as previously described with elution fractions collected that showed >90% purity (**Supplementary Figure 3**). The yield of T7 RNAP from the 1-L bioreactor showed equivalence to the petri dish, but with improved purity of 95% compared to 90% (**Figure 4B**). Finally, as previously observed both the petri dish and the bioreactor produced T7 RNAP showed improved activity compared to a commercial product (**Figure 4C**). Both produced products showed ∼196% activity compared to the commercial T7 RNAP and showed similar activity profiles to each other. These results show that T7 RNAP produced by CFE had higher activity than T7 RNAP produced by cells, which may be attractive as efforts are underway to create more active T7 RNAP for manufacturing our manufacturing pipeline could benefit production (Dousis et al., 2023).

**Figure 5.**
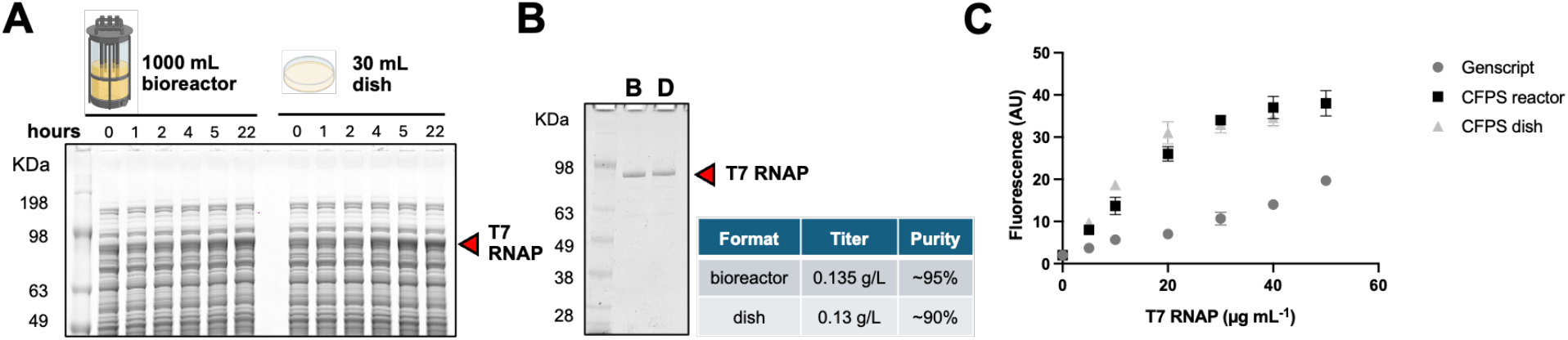
Scaled expression, purification, and activity of T7 RNA polymerase. A) Coomassie staining of an SDS-PAGE depicting the T7 RNAP CFE time course in bioreactor and “thin film” formats. The red arrow designates the band of protein accumulation over time. B) Strep-tagged T7 RNAP from either bioreactor or petri dish formats is purified and analyzed via Coomassie-staining of an SDS-PAGE. Titers are listed as per liter of CFE reaction. C) Fluorescence based T7 RNAP activity assay is used to compare bioreactor- and dish-derived polymerase to the commercially available standard (Genscript). Data are representative of n=3 independent experiments, with standard deviation shown.

## Conclusion

While CFE has been celebrated for its ability to produce proteins at speed, this is often done at the microliter scale. In this work, we have shown the power of CFE to rapidly produce T7 RNAP at the 1-L scale. Notably, reaction times were equivalent to microliter scale, and there was no drop in yield across multiple orders of magnitude in reaction volume. Our demonstration supports the power of CFE to be used for manufacturing, helping to lay the groundwork for the creation of new logistical paradigms for decentralized manufacturing. This will allow for rapid response to urgent emergent or pandemic threats and enable dependable domestic manufacturing capacity to support the bioeconomy.

## Author contributions

W.K.K., A.S.K., and M.C.J. conceived of this project. R.R., R.A., W.C., K.S., A.K., H.C., A.L., S.R., D.P., E.K., planned experiments, prepared reagents, performed experiments and analyzed data. W.K.K. and M.C.J. supervised the project. R.A., R.R., A.K., A.S.K., and M.C.J. contributed to the writing of the manuscript.

## Data availability statement

The dataset(s) supporting the conclusions of this article is(are) included within the article and its additional file(s).

## Acknowledgments

This work is supported by the Defense Advanced Research Project Agency (DARPA) contract W911NF-23-2-0039. Any opinions, findings, and conclusion or recommendations expressed in this study are those of the authors and do not necessarily reflect the view of the DARPA or the US government.

## References

Bolgiano, B., Mawas, F., Burkin, K., Crane, D. T., Saydam, M., Rigsby, P., & Corbel, M. J. (2007). A retrospective study on the quality of Haemophilus influenzae type b vaccines used in the UK between 1996 and 2004. Hum Vaccin, 3(5), 176–182. 10.4161/hv.3.5.4352

Borhani, S. G., Levine, M. Z., Krumpe, L. H., Wilson, J., Henrich, C. J., O’Keefe, B. R., Lo, D. C., Sittampalam, G. S., Godfrey, A. G., Lunsford, R. D., Mangalampalli, V., Tao, D., LeClair, C. A., Thole, A. P., Frey, D., Swartz, J., & Rao, G. (2023). An approach to rapid distributed manufacturing of broad spectrum anti-viral griffithsin using cell-free systems to mitigate pandemics. N Biotechnol, 76, 13–22. 10.1016/j.nbt.2023.04.003

Brito, L. A., & Singh, M. (2011). Acceptable levels of endotoxin in vaccine formulations during preclinical research. J Pharm Sci, 100(1), 34–37. 10.1002/jps.22267

Bundy, B. C., Hunt, J. P., Jewett, M. C., Swartz, J. R., Wood, D. W., Frey, D. D., & Rao, G. (2018). Cell-free biomanufacturing. Current Opinion in Chemical Engineering, 22, 177–183. 10.1016/j.coche.2018.10.003

Carlson, E. D., Gan, R., Hodgman, C. E., & Jewett, M. C. (2012). Cell-Free Protein Synthesis: Applications Come of Age. Biotechnol Adv, 30(5), 1185–1194. 10.1016/j.biotechadv.2011.09.016

Chan, W., Theriault, T., Levy, R., & Swartz, J. (2015). Development of a Novel Virus-like Particle (VLP) Vaccine for Personalized B-Cell Lymphoma and Chronic Lymphocytic Leukemia Therapy. Blood, 126(23), 2748. 10.1182/blood.V126.23.2748.2748

Chen, X., Li, Y., Xiong, K., & Wagner, T. E. (1994). A self-initiating eukaryotic transient gene expression system based on cotransfection of bacteriophage T7 RNA polymerase and DNA vectors containing a T7 autogene. Nucleic Acids Res, 22(11), 2114–2120. 10.1093/nar/22.11.2114

Chen, X., & Lu, Y. (2021). In silico Design of Linear DNA for Robust Cell-Free Gene Expression. Front Bioeng Biotechnol, 9, 670341. 10.3389/fbioe.2021.670341

Chou, C. P. (2007). Engineering cell physiology to enhance recombinant protein production in Escherichia coli. Appl Microbiol Biotechnol, 76(3), 521–532. 10.1007/s00253-007-1039-0

Cole, S. D., Beabout, K., Turner, K. B., Smith, Z. K., Funk, V. L., Harbaugh, S. V., Liem, A. T., Roth, P. A., Geier, B. A., Emanuel, P. A., Walper, S. A., Chávez, J. L., & Lux, M. W. (2019). Quantification of Interlaboratory Cell-Free Protein Synthesis Variability. ACS Synth Biol, 8(9), 2080–2091. 10.1021/acssynbio.9b00178

Collins, M., Lau, M. B., Ma, W., Shen, A., Wang, B., Cai, S., La Russa, M., Jewett, M. C., & Qi, L. S. (2024). A frugal CRISPR kit for equitable and accessible education in gene editing and synthetic biology. Nat Commun, 15(1), 6563. 10.1038/s41467-024-50767-2

Davanloo, P., Rosenberg, A. H., Dunn, J. J., & Studier, F. W. (1984). Cloning and expression of the gene for bacteriophage T7 RNA polymerase. Proc Natl Acad Sci U S A, 81(7), 2035–2039. 10.1073/pnas.81.7.2035

Dopp, J. L., Rothstein, S. M., Mansell, T. J., & Reuel, N. F. (2019). Rapid prototyping of proteins: Mail order gene fragments to assayable proteins within 24 hours. Biotechnol Bioeng, 116(3), 667–676. 10.1002/bit.26912

Dousis, A., Ravichandran, K., Hobert, E. M., Moore, M. J., & Rabideau, A. E. (2023). An engineered T7 RNA polymerase that produces mRNA free of immunostimulatory byproducts. Nature Biotechnology, 41(4), 560–568. 10.1038/s41587-022-01525-6

Ekas, H. M., Wang, B., Silverman, A.D., Lucks, J.B., Karim, A.S., and Jewett, M.C. (2024a). An automated, cell-free workflow for transcription factor engineering. ACS Synth Biol, In press.

Ekas, H. M., Wang, B., Silverman, A.D., Lucks, J.B., Karim, A.S., and Jewett, M.C. (2024b). Engineering a PbrR-based biosensor for cell-free detection of lead at the legal limit. ACS Synthetic Biology., In press.

Gibson, D. G., Young, L., Chuang, R. Y., Venter, J. C., Hutchison, C. A., 3rd, & Smith, H. O. (2009). Enzymatic assembly of DNA molecules up to several hundred kilobases. Nat Methods, 6(5), 343–345. 10.1038/nmeth.1318

Grodberg, J., & Dunn, J. J. (1988). ompT encodes the Escherichia coli outer membrane protease that cleaves T7 RNA polymerase during purification. J Bacteriol, 170(3), 1245–1253. 10.1128/jb.170.3.1245-1253.1988

Guan, D., Wang, D., Hallegatte, S., Davis, S. J., Huo, J., Li, S., Bai, Y., Lei, T., Xue, Q., Coffman, D. M., Cheng, D., Chen, P., Liang, X., Xu, B., Lu, X., Wang, S., Hubacek, K., & Gong, P. (2020). Global supply-chain effects of COVID-19 control measures. Nature Human Behaviour, 4(6), 577–587. 10.1038/s41562-020-0896-8

Hadi, T., Nozzi, N., Melby, J. O., Gao, W., Fuerst, D. E., & Kvam, E. (2020). Rolling circle amplification of synthetic DNA accelerates biocatalytic determination of enzyme activity relative to conventional methods. Sci Rep, 10(1), 10279. 10.1038/s41598-020-67307-9

Huang, A., Nguyen, P. Q., Stark, J. C., Takahashi, M. K., Donghia, N., Ferrante, T., Dy, A. J., Hsu, K. J., Dubner, R. S., Pardee, K., Jewett, M. C., & Collins, J. J. (2018). BioBits™ Explorer: A modular synthetic biology education kit. Sci Adv, 4(8), eaat5105. 10.1126/sciadv.aat5105

Hunt, A. C., Rasor, B.J., Seki, K., Ekas, H.M., Warfel, K.F., Karim, A.S., and Jewett, M.C. (2024). Cell-free gene expression: methods and applications. Chemical Reviews., In press.

Hunt, A. C., Vögeli, B., Hassan, A. O., Guerrero, L., Kightlinger, W., Yoesep, D. J., Krüger, A., DeWinter, M., Diamond, M. S., Karim, A. S., & Jewett, M. C. (2023). A rapid cell-free expression and screening platform for antibody discovery. Nat Commun, 14(1), 3897. 10.1038/s41467-023-38965-w

Jackson, N. A. C., Kester, K. E., Casimiro, D., Gurunathan, S., & DeRosa, F. (2020). The promise of mRNA vaccines: a biotech and industrial perspective. npj Vaccines, 5(1), 11. 10.1038/s41541-020-0159-8

Jew, K., Smith, P. E. J., So, B., Kasman, J., Oza, J. P., & Black, M. W. (2022). Characterizing and Improving pET Vectors for Cell-free Expression. Front Bioeng Biotechnol, 10, 895069. 10.3389/fbioe.2022.895069

Jewett, M. C., Calhoun, K. A., Voloshin, A., Wuu, J. J., & Swartz, J. R. (2008). An integrated cell-free metabolic platform for protein production and synthetic biology. Molecular Systems Biology, 4(1), 220. 10.1038/msb.2008.57

Jin, X., Kightlinger, W., & Hong, S. H. (2019). Optimizing Cell-Free Protein Synthesis for Increased Yield and Activity of Colicins. Methods and Protocols, 2(2), 28. https://www.mdpi.com/2409-9279/2/2/28

Karim, A. S., Dudley, Q. M., Juminaga, A., Yuan, Y., Crowe, S. A., Heggestad, J. T., Garg, S., Abdalla, T., Grubbe, W. S., Rasor, B. J., Coar, D. N., Torculas, M., Krein, M., Liew, F., Quattlebaum, A., Jensen, R. O., Stuart, J. A., Simpson, S. D., Köpke, M., & Jewett, M. C. (2020). In vitro prototyping and rapid optimization of biosynthetic enzymes for cell design. Nat Chem Biol, 16(8), 912–919. 10.1038/s41589-020-0559-0

Karim, A. S., & Jewett, M. C. (2016). A cell-free framework for rapid biosynthetic pathway prototyping and enzyme discovery. Metab Eng, 36, 116–126. 10.1016/j.ymben.2016.03.002

Kightlinger, W., Duncker, K. E., Ramesh, A., Thames, A. H., Natarajan, A., Stark, J. C., Yang, A., Lin, L., Mrksich, M., DeLisa, M. P., & Jewett, M. C. (2019). A cell-free biosynthesis platform for modular construction of protein glycosylation pathways. Nat Commun, 10(1), 5404. 10.1038/s41467-019-12024-9

Kightlinger, W., Lin, L., Rosztoczy, M., Li, W., DeLisa, M. P., Mrksich, M., & Jewett, M. C. (2018). Design of glycosylation sites by rapid synthesis and analysis of glycosyltransferases. Nat Chem Biol. 10.1038/s41589-018-0051-2

Krüger, A., Mueller, A. P., Rybnicky, G. A., Engle, N. L., Yang, Z. K., Tschaplinski, T. J., Simpson, S. D., Köpke, M., & Jewett, M. C. (2020). Development of a clostridia-based cell-free system for prototyping genetic parts and metabolic pathways. Metab Eng, 62, 95–105. 10.1016/j.ymben.2020.06.004

Kumar, G., & Chernaya, G. (2009). Cell-free protein synthesis using multiply-primed rolling circle amplification products. Biotechniques, 47(1), 637–639. 10.2144/000113171

Lee, S. J., & Kim, D. M. (2024). Cell-Free Synthesis: Expediting Biomanufacturing of Chemical and Biological Molecules. Molecules, 29(8). 10.3390/molecules29081878

Liew, F. E., Nogle, R., Abdalla, T., Rasor, B. J., Canter, C., Jensen, R. O., Wang, L., Strutz, J., Chirania, P., De Tissera, S., Mueller, A. P., Ruan, Z., Gao, A., Tran, L., Engle, N. L., Bromley, J. C., Daniell, J., Conrado, R., Tschaplinski, T. J., … Köpke, M. (2022). Carbon-negative production of acetone and isopropanol by gas fermentation at industrial pilot scale. Nature Biotechnology, 40(3), 335–344. 10.1038/s41587-021-01195-w

Lizardi, P. M., Huang, X., Zhu, Z., Bray-Ward, P., Thomas, D. C., & Ward, D. C. (1998). Mutation detection and single-molecule counting using isothermal rolling-circle amplification. Nature Genetics, 19(3), 225–232. 10.1038/898

Mao, H. H., & Chao, S. (2020). Advances in Vaccines. In A. C. Silva, J. N. Moreira, J. M. S. Lobo, & H. Almeida (Eds.), Current Applications of Pharmaceutical Biotechnology (pp. 155–188). Springer International Publishing. 10.1007/10_2019_107

McSweeney, M. A., & Styczynski, M. P. (2021). Effective Use of Linear DNA in Cell-Free Expression Systems [Review]. Front Bioeng Biotechnol, 9. 10.3389/fbioe.2021.715328

Muller, D. K., Martin, C. T., & Coleman, J. E. (1988). Processivity of proteolytically modified forms of T7 RNA polymerase. Biochemistry, 27(15), 5763–5771. 10.1021/bi00415a055

Nguyen, P. Q., Soenksen, L. R., Donghia, N. M., Angenent-Mari, N. M., de Puig, H., Huang, A., Lee, R., Slomovic, S., Galbersanini, T., Lansberry, G., Sallum, H. M., Zhao, E. M., Niemi, J. B., & Collins, J. J. (2021). Wearable materials with embedded synthetic biology sensors for biomolecule detection. Nature Biotechnology, 39(11), 1366–1374. 10.1038/s41587-021-00950-3

Pardee, K., Slomovic, S., Nguyen, P. Q., Lee, J. W., Donghia, N., Burrill, D., Ferrante, T., McSorley, F. R., Furuta, Y., Vernet, A., Lewandowski, M., Boddy, C. N., Joshi, N. S., & Collins, J. J. (2016). Portable, On-Demand Biomolecular Manufacturing. Cell, 167(1), 248–259 e212. 10.1016/j.cell.2016.09.013

Peruzzi, J. A., Vu, T. Q., Gunnels, T. F., & Kamat, N. P. (2023). Rapid Generation of Therapeutic Nanoparticles Using Cell-Free Expression Systems. Small Methods, 7(12), e2201718. 10.1002/smtd.202201718

Qin, S., Tang, X., Chen, Y., Chen, K., Fan, N., Xiao, W., Zheng, Q., Li, G., Teng, Y., Wu, M., & Song, X. (2022). mRNA-based therapeutics: powerful and versatile tools to combat diseases. Signal Transduction and Targeted Therapy, 7(1), 166. 10.1038/s41392-022-01007-w

Raghuwanshi, N., Nagar, N., Singh, S., Kaviraj, S., & Raghuwanshi, A. (2024). Purification of T7 RNA polymerase for large scale production of mRNA vaccines and therapeutics. Process Biochemistry, 147, 391–401. 10.1016/j.procbio.2024.10.005

Ramm, K., & Plückthun, A. (2000). The Periplasmic Escherichia coli Peptidylprolyl cis,trans-Isomerase FkpA: II. ISOMERASE-INDEPENDENT CHAPERONE ACTIVITY IN VITRO *. Journal of Biological Chemistry, 275(22), 17106–17113. 10.1074/jbc.M910234199

Rosa, S. S., Prazeres, D. M. F., Azevedo, A. M., & Marques, M. P. C. (2021). mRNA vaccines manufacturing: Challenges and bottlenecks. Vaccine, 39(16), 2190–2200. 10.1016/j.vaccine.2021.03.038

Sadat Mousavi, P., Smith, S. J., Chen, J. B., Karlikow, M., Tinafar, A., Robinson, C., Liu, W., Ma, D., Green, A. A., Kelley, S. O., & Pardee, K. (2020). A multiplexed, electrochemical interface for gene-circuit-based sensors. Nature Chemistry, 12(1), 48–55. 10.1038/s41557-019-0366-y

Silverman, A. D., Akova, U., Alam, K. K., Jewett, M. C., & Lucks, J. B. (2020). Design and Optimization of a Cell-Free Atrazine Biosensor. ACS Synth Biol, 9(3), 671–677. 10.1021/acssynbio.9b00388

Silverman, A. D., Karim, A. S., & Jewett, M. C. (2020). Cell-free gene expression: an expanded repertoire of applications. Nat Rev Genet, 21(3), 151–170. 10.1038/s41576-019-0186-3

Spice, A. J., Aw, R., Bracewell, D. G., & Polizzi, K. M. (2020). Synthesis and Assembly of Hepatitis B Virus-Like Particles in a Pichia pastoris Cell-Free System. Front Bioeng Biotechnol, 8, 72–72. 10.3389/fbioe.2020.00072

Stark, J. C., Huang, A., Hsu, K. J., Dubner, R. S., Forbrook, J., Marshalla, S., Rodriguez, F., Washington, M., Rybnicky, G. A., Nguyen, P. Q., Hasselbacher, B., Jabri, R., Kamran, R., Koralewski, V., Wightkin, W., Martinez, T., & Jewett, M. C. (2019). BioBits Health: Classroom Activities Exploring Engineering, Biology, and Human Health with Fluorescent Readouts. ACS Synth Biol, 8(5), 1001–1009. 10.1021/acssynbio.8b00381

Stark, J. C., Huang, A., Nguyen, P. Q., Dubner, R. S., Hsu, K. J., Ferrante, T. C., Anderson, M., Kanapskyte, A., Mucha, Q., Packett, J. S., Patel, P., Patel, R., Qaq, D., Zondor, T., Burke, J., Martinez, T., Miller-Berry, A., Puppala, A., Reichert, K., … Jewett, M. C. (2018). BioBits™ Bright: A fluorescent synthetic biology education kit. Sci Adv, 4(8), eaat5107. 10.1126/sciadv.aat5107

Stark, J. C., Jaroentomeechai, T., Moeller, T. D., Hershewe, J. M., Warfel, K. F., Moricz, B. S., Martini, A. M., Dubner, R. S., Hsu, K. J., Stevenson, T. C., Jones, B. D., DeLisa, M. P., & Jewett, M. C. (2021). On-demand biomanufacturing of protective conjugate vaccines. Sci Adv, 7(6), eabe9444. 10.1126/sciadv.abe9444

Stark, J. C., Jaroentomeechai, T., Warfel, K. F., Hershewe, J. M., DeLisa, M. P., & Jewett, M. C. (2023). Rapid biosynthesis of glycoprotein therapeutics and vaccines from freeze-dried bacterial cell lysates. Nat Protoc, 18(7), 2374–2398. 10.1038/s41596-022-00799-z

Sun, Z. Z., Yeung, E., Hayes, C. A., Noireaux, V., & Murray, R. M. (2014). Linear DNA for Rapid Prototyping of Synthetic Biological Circuits in an Escherichia coli Based TX-TL Cell-Free System. ACS Synth Biol, 3(6), 387–397. 10.1021/sb400131a

Tabor, S., & Richardson, C. C. (1985). A bacteriophage T7 RNA polymerase/promoter system for controlled exclusive expression of specific genes. Proc Natl Acad Sci U S A, 82(4), 1074–1078. 10.1073/pnas.82.4.1074

Thavarajah, W., Silverman, A. D., Verosloff, M. S., Kelley-Loughnane, N., Jewett, M. C., & Lucks, J. B. (2020). Point-of-Use Detection of Environmental Fluoride via a Cell-Free Riboswitch-Based Biosensor. ACS Synth Biol, 9(1), 10–18. 10.1021/acssynbio.9b00347

Vögeli, B., Schulz, L., Garg, S., Tarasava, K., Clomburg, J. M., Lee, S. H., Gonnot, A., Moully, E. H., Kimmel, B. R., Tran, L., Zeleznik, H., Brown, S. D., Simpson, S. D., Mrksich, M., Karim, A. S., Gonzalez, R., Köpke, M., & Jewett, M. C. (2022). Cell-free prototyping enables implementation of optimized reverse β-oxidation pathways in heterotrophic and autotrophic bacteria. Nat Commun, 13(1), 3058. 10.1038/s41467-022-30571-6

Warfel, K. F., Williams, A., Wong, D. A., Sobol, S. E., Desai, P., Li, J., Chang, Y.-F., DeLisa, M. P., Karim, A. S., & Jewett, M. C. (2023). A Low-Cost, Thermostable, Cell-Free Protein Synthesis Platform for On-Demand Production of Conjugate Vaccines. ACS Synth Biol, 12(1), 95–107. 10.1021/acssynbio.2c00392

Williams, A. J., Warfel, K. F., Desai, P., Li, J., Lee, J. J., Wong, D. A., Nguyen, P. M., Qin, Y., Sobol, S. E., Jewett, M. C., Chang, Y. F., & DeLisa, M. P. (2023). A low-cost recombinant glycoconjugate vaccine confers immunogenicity and protection against enterotoxigenic Escherichia coli infections in mice. Front Mol Biosci, 10, 1085887. 10.3389/fmolb.2023.1085887

Yin, G., Garces, E. D., Yang, J., Zhang, J., Tran, C., Steiner, A. R., Roos, C., Bajad, S., Hudak, S., Penta, K., Zawada, J., Pollitt, S., & Murray, C. J. (2012). Aglycosylated antibodies and antibody fragments produced in a scalable in vitro transcription-translation system. mAbs, 4(2), 217–225. 10.4161/mabs.4.2.19202

Zawada, J. F., Yin, G., Steiner, A. R., Yang, J., Naresh, A., Roy, S. M., Gold, D. S., Heinsohn, H. G., & Murray, C. J. (2011). Microscale to manufacturing scale-up of cell-free cytokine production—a new approach for shortening protein production development timelines. Biotechnol Bioeng, 108(7), 1570–1578. 10.1002/bit.23103

